# MODE for detecting and estimating genetic causal variants

**DOI:** 10.1101/357228

**Authors:** V. S. Sundar, Chun-Chieh Fan, Dominic Holland, Anders M. Dale

## Abstract

Determining the genetic causal variants and estimating their effect sizes are considered to be correlated but independent problems. Fine-mapping studies often rely on the ability to integrate useful functional annotation information into genome wide association univariate/multivariate analysis. In the present study, by modeling the probability of a SNP being causal and its effect size as a set of correlated Gaussian/non-Gaussian random variables, we design an optimization routine for simultaneous fine-mapping and effect size estimation. The algorithm is released as an open source C package MODE.

**Availability and Implementation:** http://sites.google.com/site/sundarvelkur/mode

## 1 Introduction

Detection and estimation of the genetic causal variants associated with a particular phenotypic trait is typically accomplished by reinforcing Genome Wide Association Studies (GWAS) findings with fine mapping analysis. However, for highly polygenic phenotypes, more often than not, biologically causal SNPs do not reach genome-wide significance [1, 2, 3, 4, 5, 6]. In this work, we aim to simultaneously estimate the probability of a SNP being causal and its effect size by developing a well designed optimization routine, that allows for incorporation of functional annotation data. This could potentially aid in accurate detection and estimation of causal loci.

## 2 Problem statement

Modeling the genotype-phenotype relation through a linear model [7]

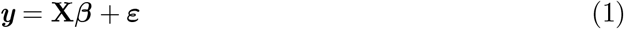

where *N* is the number of subjects, *n* is the number of genetic markers, ***y*** is a *N* × 1 phenotype vector, **X** related is the *N × n* genotype matrix, and *ɛ* is a *N* × 1 is the vector of noise terms modeled as *N*(0, Σ_*ε*_), we aim to estimate the regression coefficients *β* such that 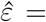 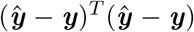 is minimum. The elements of **X** are typically coded as 0,1 or 2. It is well documented that due to the correlated and sparse nature of the SNPs, the univariate regression results in erroneous estimates [8, 9, 10]. Following [11], we minimize *F* = *L* + *C*, with

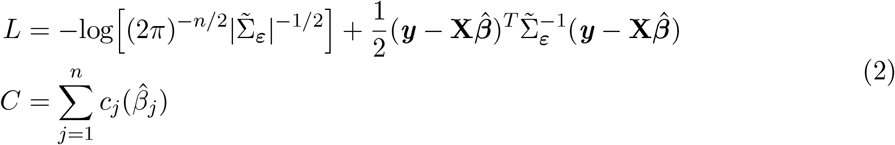

where the cost associated with the *j*^th^ SNP is given as

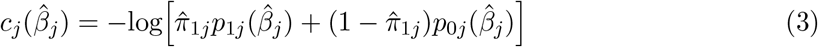

Here 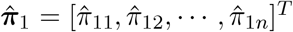 is the *n* × 1 vector of non-null prior probabilities of the SNPs. *p*_1*j*_ (•) and *p*_0*j*_ (•) denote the pdf of causal and null SNPs, respectively. The causal variants and their effect sizes are obtained by minimizing *F* with respect to *π*_1_ and *β*. Due to Linkage Disequilibrium (LD) and other covariates, the effect sizes and the prior probabilities could be correlation. We take into account these correlations while solving for *π*_1_ and *β*. Minimization of the two-term objective function (likelihood and cost functions) is carried out efficiently using the conjugate gradient method.

## 3 Method

The function to be minimized is a combination of an error minimizing term - the negative log-likelihood function, and a cost term which imparts the necessary sparse characteristics to the effect sizes. We model the probability of a SNP being causal as uniformly distributed between *a* and *b*, i.e. *π*_1*j*_ ~ *U*[*a*_*j*_,*b*_*j*_], and the effect sizes as a Gaussian distribution, *β*_*j*_ ~ *N*(*μ*, *σ*). Probabilistic shrinkage is achieved by modeling the null SNPs using a Laplace pdf with zero mean and *σ*_0_ standard deviation. Incorporation of Linkage Disequilibrium and function annotation information is through the distributions of prior probabilities, effect sizes, and the correlations among them. The 2*n* × 2*n* correlation matrix structure is given as

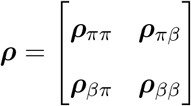

If no information about the correlation structure is known, the ***ρ*** matrix is taken to be the identity matrix. The correlated non-Gaussian random variables are first transformed into standard normal random variables using the Nataf’s transformation [12], as implemented in [13]. Central difference scheme is used to obtain the gradients of the cost function with respect to *β* and *π*_1_. Mathematical details regarding the gradients and hessian (with respect to *β*) can be found in [11]. A similar approach can be used in obtaining the derivatives with respect to *π*_1_. MODE source code and binary executables can be downloaded from http://sites.google.com/sites/sundarvelkur/mode. MODE borrows functions from ART [13], an open source package for simulation of correlated non-Gaussian random variables.

## 4 Results

### 4.1 Simulation studies

The phenotype vector with a heritability 0.5 is simulated for 100000 individuals using Eq. (1) utilizing the genotype matrix obtained using Hapgen2 [14] and 1000 Genomes [15]. We consider the first 20000 SNPs of chromosomes 1 to 22 with minor allele frequency greater than 0.01. The number of causal variants are taken to be 50% of all the SNPs belonging to the functional annotation {Exon, 3’UTR, 5’UTR}, with effect sizes distributed as *N*(0, 1). We consider three replication sets and three different effect size vectors, resulting in 18 cases to estimate the mean positive predictive value (PPV), negative predictive value (NPV), and correlation between the estimated and true effect size. MODE results, along with few other techniques [11] are shown in Figure 1.

**Figure 1:**
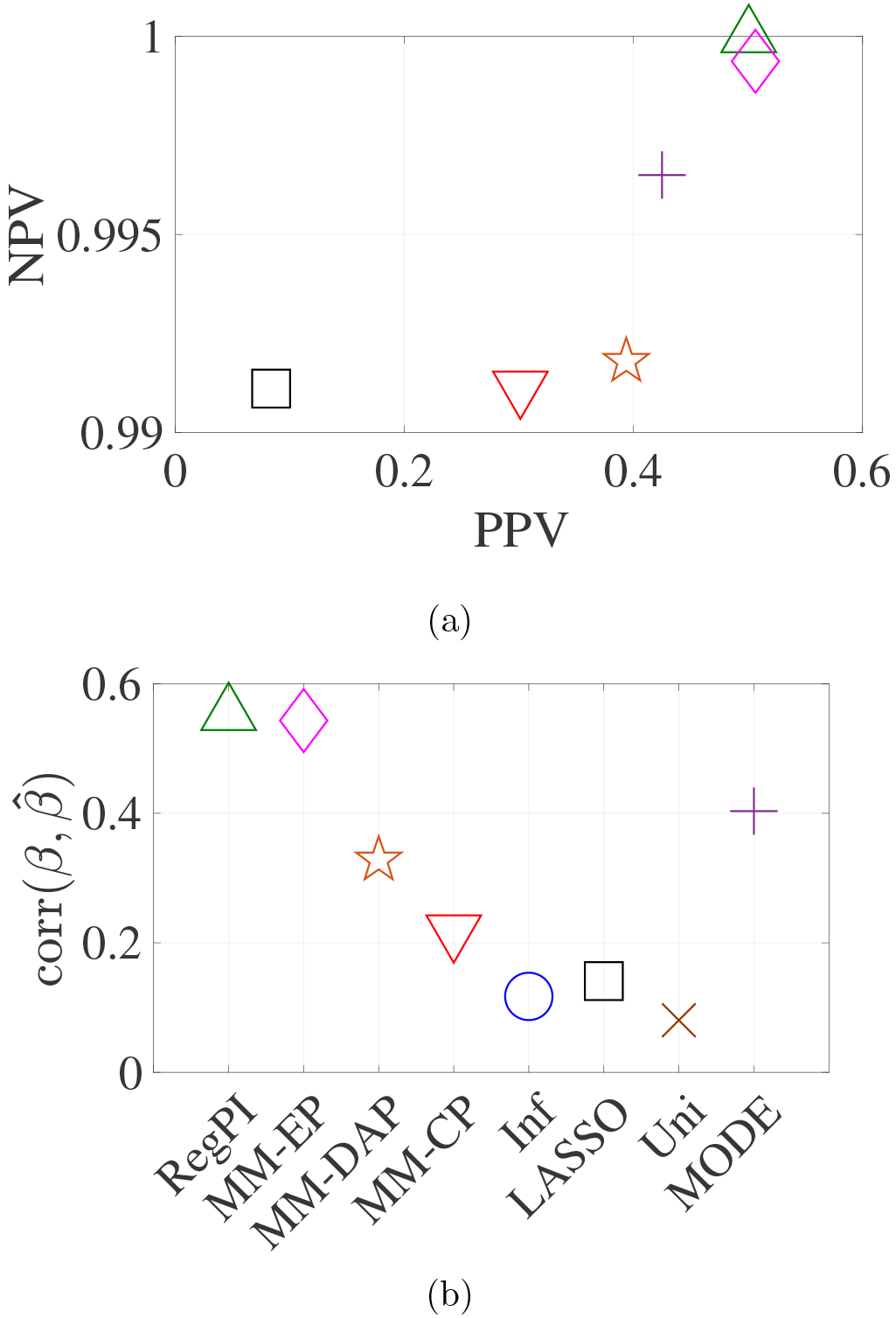
(a) correlation between the estimated and true effect sizes, (b) PPV and NPV. RegPI: Regularized pseudo inverse (green triangle); MM-EP: Mixture model with enriched priors (magenta diamond); MM-DAP: Mixture model with DAP priors (orange star); MM-CP: Mixture model with constant priors (red inverted triangle); Infinitesimal: Normal prior (no mixture) (blue circle); LASSO (black square); Univariate (brown cross); MODE (purple plus). Refer to [11] for the details of the methods.

## 5 Discussions

It is observed that in comparison with the MM-CP and MM-DAP [6], MODE estimates have better PPV and NPV characteristics. The improvement in the correlation is due to better NPV of MODE. The relationship between the causal nature of the SNPs and its effect size, and correlations among the causal probabilities and effect sizes are typically unknown. Specification of these quantities require heuristics or additional information, possibly from gene-network analysis, which could identify potential causal SNPs and their relationship with adjacent SNPs. The algorithm is computational tractable unlike MCMC based bayesian methods which requires heavy computational resources and time. MODE locates the causal SNPs and estimates its effect size efficiently with acceptable accuracy.

